# The IgE production is initially induced in subcutaneous fat and depends on extrafollicular B cells

**DOI:** 10.1101/2021.08.18.456888

**Authors:** Dmitrii Borisovich Chudakov, Gulnar Vaisovna Fattakhova, Mariya Vladimirovna Konovalova, Daria Sergeevna Tsaregorotseva, Marina Alexandrovna Shevchenko, Olga Dmitrievna Kotsareva, Anton Andreevich Sergeev, Elena Victorovna Svrshcehvskaya

**Affiliations:** Laboratory of Cell Interactions, Shemyakin-Ovchinnikov Institute of Bioorganic Chemistry, Moscow, Russian Federation; Sechenov First Moscow State Medical University, Moscow, Russian Federation.

**Keywords:** IgE, Extrafollicular response, subcutaneous fat, plasmablasts, T-extrafollicular helpers.

## Abstract

**Background:** Growing body of evidence indicates that IgE production can be developed by mechanisms that differ from those responsible for IgG and IgA production. One potential possibility is generation of IgE producing cells from tissue-associated B-cells and/or through extrafollicular pathway. But the role of subcutaneous fat-associated B-cells in this process is poorly investigated. The aim of the present study was to investigate the role of different B- and T-cell subpopulations after long-term antigen administration in IgE response.

**Methods:** BALB/c mice were immunized 3 times a weeks for 4 weeks in withers region enriched with subcutaneous fat with high and low antigen doses as well as by intraperitoneal route in region enriched with visceral fat for comparison.

**Results:** After long-term antigen administration that promotes the type of immune response which is more similar to one observed in young allergic children, subcutaneous fat tissue B-cells generates more rapid and active IgE class switched and IgE-produced cells. Although IgE production at later time points was initiated also in regional lymph nodes, the early IgE production was exclusively linked with subcutaneous fat. We observed that low-dose induced strong IgE production accompanied by minimal IgG1 production was linked with extrafollicular B-2 derived plasmablasts as well as extrafollicular T- helpers accumulation. Delayed IgE class switching in regional lymph nodes and visceral fat tissue was characterized by the absence of both stable plasmablasts and T-extrafollicular helpers accumulation.

**Conclusion:** Extrafollicular B- and T-cell responses in subcutaneous fat are necessary for early IgE class switching and sensitization process in the case of allergen penetration through skin.

## Introduction

It is generally accepted that the predisposition to development of allergic immune response in individuals depends on their barrier tissue properties, including lymphoid structures located in the vicinity to these barrier tissues. For example, polymorphisms of *FLG* gene encoding fillagrin protein associated with skin barrier function as well as in *IL33* and *TSLP* genes encoding cytokines upregulated in response to barrier tissue damage are linked with atopic dermatitis development [1]. Some genetic variants of claudin-1 gene and different expression levels of claudin-18 gene is linked with atopic dermatitis and asthma development, respectively [2, 3]. These features in gene structure and/or expression levels may be responsible for poor tight junctions formation and for predisposition to barrier tissues damage [1] which, in turn, leads to production of type 2 response inducing cytokines and subsequent T-helper cell polarization and IgE production [4]. Nowadays the important role of local tissue-associated tertiary lymphoid structures in initial studies of allergic immune response development becomes more evident. From clinical studies of nasal polyps in patients with allergic rhinitis [5; 6], as well as some experimental mouse asthma models [7–9], it becomes clear that mucosal-associated B-cells could be activated to class switch recombination *in situ*, in nasal polyps or inducible bronchial-associated lymphoid tissue (iBALT). It is also interesting that at least at some conditions this process can be dependent more strikingly from extrafollicular B-cell activation and requires minimal GC response [10].

Although allergens could penetrate not only via airway epithelium but also through skin epithelium with subcutaneous adipose tissue, the role of fat-associated lymphoid clusters (FALCs) or at least fat-associated B-cells in the initial studies of allergic immune response is not well studied in comparison to the role of nasal polyps and iBALT. Although it is believed that in subcutaneous fat the number of FALCs is more lower than in abdominal fat or epicardium [11] and that visceral fat-associated B-cells are linked with chronic inflammation for example during the development of insulin resistance [12], the other studies indicate that subcutaneous fat, especially in obese individuals, contains immunologically active B- and T-cells [13; 14]. It is important to mention that obesity is one of the lifestyle factors linked with asthma development [15] and administration of pro-adipogenic leptin enhances IgE production and asthma development in mice [16] while administration of anti-adipogenic adiponectin inhibited it [17]. These facts indicate that subcutaneous fat tissue is mainly linked with obesity, and fat-associated B-cells can be responsible for initiation of pro-allergic immune response especially when low antigen doses penetrates skin barrier to enter the internal environment of the body.

We and others [18–20] have shown that specific IgE production in allergic patients is not linked with specific IgG1 or IgG4 production at least in the case of non-replicative allergens. This fact indicates that specific IgE production is induced by mechanism different from that responsible for IgG production. It is well known that high IgG response requires strong GC induction, mostly in secondary lymphoid organs. So, one could suppose that IgE production is triggered in the site different from secondary lymphoid organ B-cell follicles and without significant GC induction.

In most currently used allergic models, high doses of antigens are administrated to mice together with adjuvants [21–23] which induced robust GC formation. In contrast, adjuvant-free low antigen doses models can establish significant pro-anaphylactic IgE-based immune response [24–26] and that low-dose IgE inducing strategy better reflects the natural sensitization process [27]. In our previously work, we have shown that low rather than high, chronically administrated antigen doses induce significant IgE response with minimal IgG production [28]. In addition, we have shown that the route of antigen administration has significant effect on the intensity of specific IgE production. Administration of OVA as a model antigen in withers adipose tissue by subcutaneous route which resembles antigen penetration through damaged skin and entering into the subcutaneous fat induced B-cell IgE isotype switching in adipose tissue but not in regional lymph nodes. It should be mentioned that withers region in mice contains the most developed subcutaneous fat structures [29]. In the same time, administration of antigen by intraperitoneal route in the region enriched in visceral fat that, however, is not linked with barrier tissues, directly induces weak, if any, specific IgE production [28]. The mechanisms which mediate participation of B and T-cell populations in such response, as well as more delayed IgE class switching in regional lymph nodes and abdominal fat tissues are not fully understood.

In many allergic and asthma models, B-cell and T-cell subpopulations were thoroughly studied. But most authors focused on late phases of allergy immune response which are characterized by intensive manifestations of asthma or allergy symptoms and develop several days after intensive challenge starting [30]. This approach, however, provides little information on early stages of allergy development observed in children [20] before development of atopic march in later years [31], and compensatory IgG production [32] obviously related to classical GC response in some individuals. So, in this work, we focus on investigation of early stages of allergen-specific immune response.

The aim of the present work is to estimate B-cell subpopulations responsible for early IgE B-cell class switching in subcutaneous fat in comparison to regional lymph nodes, role of certain T-cell subpopulations, as well as to understand potential mechanisms of IgE class switching hampering in regional lymph nodes and abdominal fat tissue. Deeper understanding of these mechanisms will help to improve current strategies for allergy and asthma prevention in predisposed individuals.

## Materials and methods

### Mice

All animal experiment protocols were carried out by IBCh RAS IACUUC protocol. Female BALB/c mice (6-8 weeks) were obtained from Andreevka Center (Stolbovaya, Russian Federation). Mice were housed 2 weeks in SPF condition before each experiment. During this time as well as experimental protocol animals were kept in 12-h light-dark circle at room temperature in plastic cages (10-12 mice per cage). Mice were fed *ad libitum*.

### Immunization, allergen challenge and sample collection

Mice received OVA (Sigma Aldrich, Darmstadt, Germany) as a model antigen 3 times a week for 4 weeks (28 days) (Figure S1). OVA was administrated by subcutaneous route (s.c.) in withers region (W) in low (100 ng) or high (10 µg) dose or by intraperitoneal route (i.p.) in low dose. Antigen was administrated in sterilized saline solution in 100 µl volume. Intact mice or saline-treated animals were used as control groups. There were 20 mice in each experimental group. Every 7 days 5 mice from each three experimental group were challenged with 0.2 ml of 0.25% OVA solution to estimate anaphylaxis severity. Body temperature was measured by infrared thermometer CEM DT-8806S (SEM Test Instruments, Moscow, Russia) as it was performed in [33]. The temperature was measured every 15 minutes for 1.5 hour. We observed that the most significant temperature decline was detected after 45 minutes, and the magnitude of this decline was considered as a quantitative indicator of anaphylaxis severity. The magnitude of this decline is always was not higher than 2.5 °C in the case of animal survival. In some cases, however, we observed animals’ death after 30-60 minutes upon challenge. In lethal cases, anaphylaxis severity is believed to be higher than in survival cases, and the earlier death means the higher severity. Therefore, we assigned the value of –dT «3» to death time point 1 hr, value «4» to death time point 45 minutes, and value «5» to death time point 30 minutes after upon challenge.

After systematic anaphylaxis intensity measurement, mice were bled. The blood was taken by retroorbital technique from living anesthesized animals and by cardiac puncture *post mortem*. Serums were collected by centrifugation and store at -20°C. Mice were sacrificed by isoflurane («Aeran», Baxter) inhalation and perfusion through retroorbital sinus was performed. Withers adipose tissue samples or abdominal adipose tissue and regional lymph nodes were collected. For quantitative PCR samples from adipose tissue were homogenized in ExtractRNA (Evrogen) which is Trizol analog. For flow cytometry homogenization was performed in PBS pH=7.2. Homogenates were than centrifugeted (300 g) and washed 2 times with PBS. Regional lymph nodes were initially homogenized in PBS following centrifugation. 5*10^5^ cells were taken from suspension, pelleted and resuspended in ExtractRNA for gene expression levels measurement. The remaining cells were taken for flow cytometry.

### ELISA for OVA specific antibody assay

ELISA for detection of specific IgE production was carried out on 96-well microtitre plates (Costar, Maxisorb) coated with 50 μl of 20 µg/ml OVA solution in PBS pH=7.2 overnight at +4°C. After extensive washing with PBS with 0.05% Tween-20 (PBS-T) and subsequent blocking with 5% BSA in PBS, incubation with different serum dilutions was performed at +4°C overnight. Incubation with with anti-mouse IgE - HRP (clone 23G3, Abcam, 1:1000 dilution, 3 hrs) was performed next day. Plates was further processed with 3,3’5,5’-tetramethylbenzidine (TMB) substrate. Optical densities (ODs) were measured by automatic plate reader (Thermo Fisher Scientific, Waltham, MA, USA) at 450 nm with substraction of optical density at 620 nm which does not correspond to the reaction colored product. Antibody quantities were estimated as serum titers corresponded to the maximal serum dilutions where OD was three standard deviations higher than mean background OD.

The ELISA protocol for specific IgG1 production was slightly modified. Coating was performed by 5 µg/ml OVA solution in PBS pH=7.2. Blocking was performed by 1% BSA in PBS.

Following serum samples incubation, plates were further processed with anti-mouse IgG1 (clone RMG1-1, BioLegend) in 1:1000 dilution for 2 hours.

### Gene expression measurement

RNA was extracted by phenol-chloroform extraction method followed by RNAse free DNAse treatment (Thermo Fisher Scientific, USA). For measurement of DNA excision circles corresponding to direct and sequential IgE switch, we did not perform DNA digestion. cDNA was synthesized by using RevertAid First Strand cDNA Synthesis Kit (Thermo Fisher Scientific). Quantitative PCR was performed with kits from BioLabMix (Novosibirsk, Russia). Probes with 6-FAM as a fluorescent dye on 5’-end and BHQ-1 as a quencher on 3’-end were used. Expression of target genes and presence of excision DNA circles was estimated by normalizing to expression of 2 house-keeping genes, GAPDH and HPRT, and was calculated as 2^-d(dCt) in comparison with expression in the tissues of intact control mouse. Reaction was performed in CFX Connect Amplificator (BioRad) according to the following protocol: +95°C initial denaturation for 3 minutes followed by 50 cycles: 5 s denaturation at +95°C; 20 s annealing and elongation at +64°C. Reaction was performed in 96-well plates (MLP9601, BioRad) in 20 µl volume. Forward and reverse primer concentrations were 0.4 µM each, and probe concentration was 0,2 µM. Primers and probes were designed in NIH Primer BLAST and synthesis was performed by Evrogen (Moscow, Russian Federation).

The following primers and probes were used:

GAPDH:

F: GGAGAGTGTTTCCTCGTCCC; R: ACTGTGCCGTTGAATTTGCC; Z: /6-FAM/- CGCCTGGTCACCAGGGCTGCCATTTGCAGT-/BHQ-1/; product length 202 b.p.

HPRT:

F: CAGTCCCAGCGTCGTGATTA; R: TCCAGCAGGTCAGCAAAGAA; Z: /6-FAM/- TGGGAGGCCATCACATTGTGGCCCTCTGTGTG /BHQ-1/; product length 228 b.p.

germline ε

F: CCCACTTTTAGCTGAGGGCA; R: CTGGTTAAGGGCAGCTGTGA; Z: /6-FAM/-CGCCTGGGAGCCTGCACAGGGGGC-/BHQ-1/; product length 244 b.p.

circular µ-ε

F: CCCACTTTTAGCTGAGGGCA; R: CGAGGGGGAAGACATTTGGG; Z: /6-FAM/-CGCCTGGGAGCCTGCACAGGGGGC-/BHQ-1/; product length 203 b.p.

circular γ1-ε

F: AGATTCACAACGCCTGGGAG; R: GTCACTGTCACTGGCTCAGG; Z: /6-FAM/-CCACTGGCCCCTGGATCTGCTGCCCA-/BHQ-1/; product length 211 b.p.

germline γ1

F: AGAACCAAGGAAGCTGAGCC; R: AGTTTGGGCAGCAGATCCAG; Z: /6-FAM/-AGGGGAGTGGGCGGGGAGGCCA-/BHQ-1/; product length 109 b.p.

### Flow cytometry

Cells from adipose tissue and lymph node homogenates were passed through 80 µm mesh filter to obtain single cell population. Cells were washed in PBS and stained with antibodies to respective markers. To discriminate B-cell subpopulations antibodies were used as follows: CD5- BV510 (clone 53-7.3, BioLegend), CD1d-FITC (clone 1B1, BioLegend), CD95-PE (clone SA367H8, BioLegend), CD38-PECy7 (clone 90, BioLegend), CD19-APC (clone 6D5, BioLegend), B220-APCCy7 (clone RA3-6B2, BioLegend). To discriminate ILC2, NK-cells and T-cells subpopulations organs homogenates were stained: CD4-BV510 (clone GK1.5, BioLegend), CD49b-PE (clone HMa2, BioLegend), CXCR4-PerCPCy5.5 (clone L276F12, BioLegend), CXCR5-PECy7 (clone L138D7, BioLegend), ST2-APC (clone DIH4, BioLegend), CD45-APCCy7 (clone 30F11, BioLegend), biotinylated anti-lineage cocktail (Biolegend, cat # 133307) followed by streptavidin-FITC (Biolegend). For isotype control we used anti-rat IgG1 antibodies (clone GO114F17) labeled with BV510, FITC, PE, PerCPCy5.5, PECy7, APC and APCCy7 respectively.

After pre-blocking with 10% of rabbit serum for 15 minutes cells were stained for 1 hour at +4°C in FACS buffer (0,5% BSA, 0,01% NaN3 in PBS pH=7.2). To exclude death cells DAPI was added prior to flow cytometry.

B-1a cells were identified as CD19^+^B220^-^CD5^+^ [34], MZ-B cells as CD19^+^B220^+^CD1d^+^ [34], GCs as CD19^+^B220^+^CD38^-^CD95^+^ [35], extrafollicular plasmablasts as CD19^+^B220^-^ [36]. Different expression of CD38 and CD95 allows to discriminate these populations. ILC2 were identified as CD45^+^Lin^-^ST2^+^ cells [37], NK-cells as CD45^+^CD49b^+^, T-follicular helpers as CD45^+^Lin^+^CD4^+^CXCR4^+^CXCR5^+^, extrafollicular T- helpers as CD45^+^Lin^+^CD4^+^CXCR4^+^CXCR5^-^ based on the observations from [38], Th-2 as CD45^+^Lin^+^CD4^+^CXCR4^-^CXCR5^-^ST2^+^ cells [39].

For intracellular IgE staining cells were also pre-blocked with anti-mouse IgE (RME-1, BioLegend) in 1:50 final dilution for 30 minutes. After staining for surface markers, they were fixed in 4% PFA for 20 minutes and permeabilized in 0.1% Triton-X100. During permeabilization step, anti-mouse IgE-FITC (RME-1) was added.

Flow cytometry was performed on MACSQuant Tyto (Miltenyi Biotech, Germany).

Results were processed in FlowJo V10 (BD, USA).

### Histology with H&E staining

Lung tissue samples with the lower part of trachea were taken from isoflurane sacrificed mice. Lungs were immediately filled with 4% PFA through a cannula inserted into the trachea. Lungs were kept in 4% PFA before histological sections preparation. Histological sections 8 nm thick were made on microtome (???). H&E staining was performed with Hematoxylin and Eosin staining kit (Abcam, Cat # ab245880), according to manufacturer instructions.

### Statistics

All experiments were performed 2-3 times. For comparison of experimental groups, Mann-Whitney non-parametric test was used. Levels of p < 0.05 were considered statistically significant. For correlation coefficients determination, Spearman test was used. Mean and standard deviations for each compared group were calculated.

## Results

### Chronically low dose antigen administration in subcutaneous but not abdominal fat tissue induces early B-cell IgE class switching and reproduces IgE-mediated type I hypersensitivity

We have previously shown [28] that long-term antigen administration induces high specific IgE titres mainly when antigen administrated in withers region by s.c. route. Indeed, we observed that low (100 ng) dose of OVA induced substantial IgE response from 14^th^ day of immunization protocol. This response reached a plateau on 21^th^ day, as well as specific IgG1 response, when mice developed pro-anaphylactic immune response according to existence of significant temperature drop after high dose allergen challenge (Fig. 1A-C). It is interesting that in these experiment series we also observed specific IgE production in high (10 µg) dose immunized animals which was comparable to specific IgE production in low dose group to 28^th^ day (Fig. 1G). It should be mentioned that this production reached such level only at 28^th^ day and, that is more important, was accompanied by very high specific IgG_1_ production (Figure 1 D-F, H). Although anaphylactic severity in high dose immunized animals was higher than in low dose immunized mice (Fig. 1I) and allergen challenge provoked high mortality in high dose group, we did not observe significant correlation between IgE production and this anaphylactic severity (Fig. 1K). So, in this case anaphylaxis could be due to presence of pro-anaphylactic IgG1 antibodies which could trigger in some cases mast cell degranulation [40] or platelet activation factor release from macrophages [41]. Both pathways are not developed during classical human type I allergy response [42]. In contrast, we observed significant correlation between IgE production and anaphylaxis severity in low dose group (Fig. 1J). So, chronical administration of low antigen doses is characterized by high allergen specific IgE and low IgG levels, and significant dependence of anaphylaxis on specific IgE levels. Therefore, this model better reproduces clinical pathogenesis of sensitization and IgE – mediated type I allergy development in humans.

**Figure 1.**
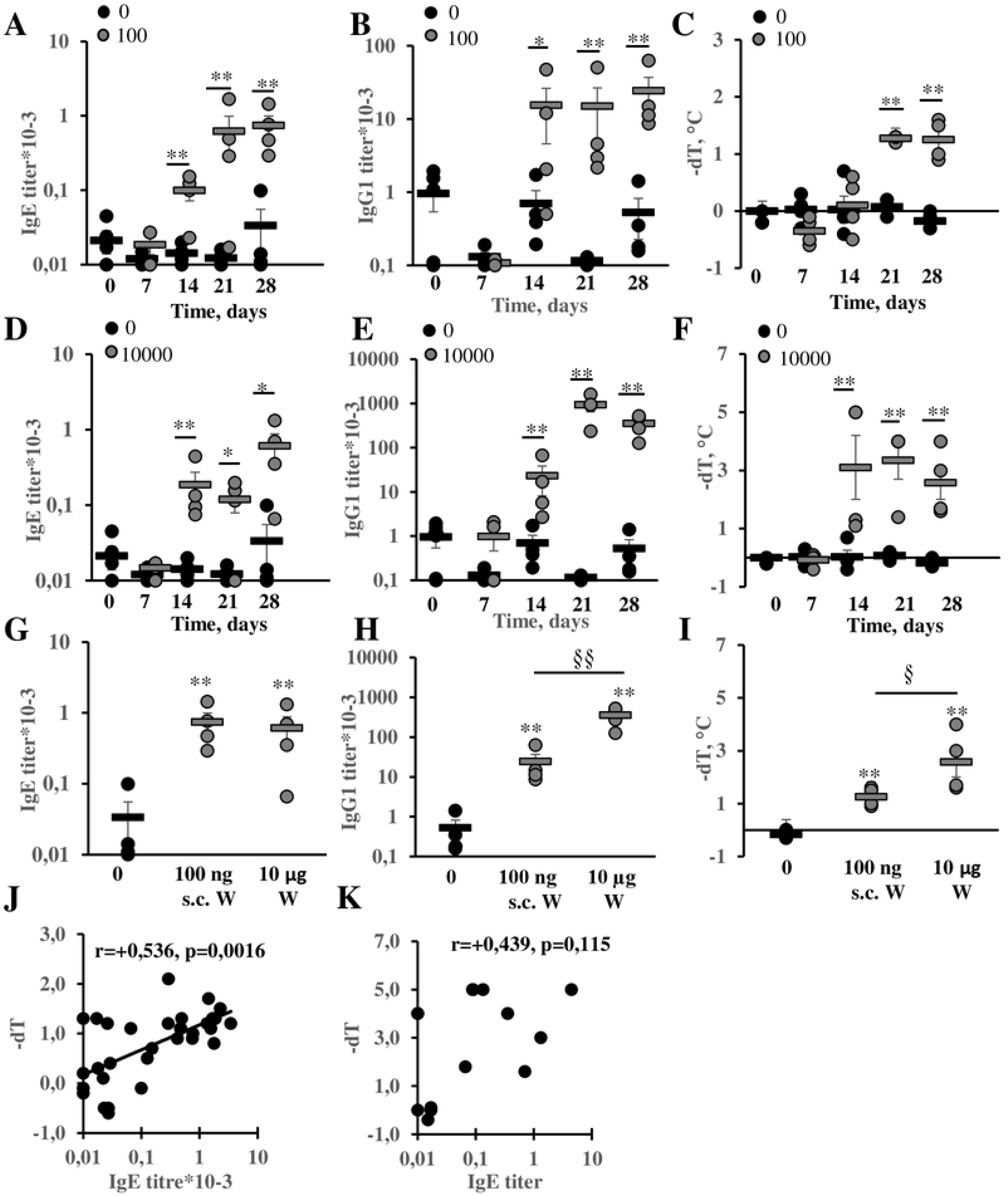
Low but not high dose immunization represents clinically relevant mouse allergy model characterized by high IgE production, anaphylaxis intensity and minimal IgG1 production. BALB/c mice were immunized by low (100 ng) (A-C) or high (10000 ng) (D-F) OVA doses 3 times a week for 4 weeks. Specific IgE (A, D) and IgG1 (B, E) titers, temperature changes (C, F) were measured at different time points upon allergen challenge and compared to saline immunized animals. Comparison of IgE (G), IgG1 (H) production and anaphylaxis severity (I) between high and low dose immunized mice at 28^th^ day. Correlations between IgE production and anaphylaxis severity in low (n=24) (J) and high dose (n=11) (K) groups. */** - p <0.05/0.01 between indicated group and saline immunized mice. §/§§ - p<0.05/0.01 between crossbars marked groups.

### Subcutaneous fat-associated B-cells are responsible for early B-cell IgE class switching

We next addressed the question if B-cell IgE class switching occurs exclusively in subcutaneous tissue associated B-cells and if these B-cells form FALCs. In our previously work [28], we have shown that B-cell IgE isotype switching, but not IgE production, was triggered in tissue associated B-cells. In this study, we used more accurate method of gene expression quantification, namely, probe based quantitative PCR instead of SybrGreen I based technique. Indeed, in accordance with our previous results, we showed that at early time points from 7^th^ to 21^th^ day of immunization induction of markers linked with isotype switching occured almost exclusively in withers adipose tissue. Meanwhile the circular µ-ε DNA excision circles were detected in lymph nodes of high dose and low dose mice group on 14^th^ day and 21^th^ day respectively (Fig. 2A-B). As far as these excision circles could originate from B-cells that had recently migrated to lymph nodes from the site of isotype switching, it is more likely that during the first 3 weeks of chronical antigen administration B-cell IgE isotype switching occured exclusively in subcutaneous fat-associated B-cells. These data were confirmed by flow cytometry of IgE^+^ B-cells quantification (Fig. 2C-D). We clearly discriminated two IgE^+^ B-cells subpopulations, namely, IgE^low^ and IgE^high^ B-cells (Fig. 2C). But despite this fact, the percentage of both IgE+ B-cell subpopulations increased firstly in adipose tissue (21^th^ day) and only secondly in lymph nodes (28^th^ day) (Fig. 2D). However, in contrast to our previous results [28], we observed expression of markers associated with B-cell IgE class switching also in regional lymph nodes after 4 weeks of antigen administration (Fig. 2A-D). It can indicate that after long time, even low antigen doses could accumulate in sufficient quantity not only in the sites of antigen administration but also in regional lymph nodes. According to another possibility, the antigen reached the lymph nodes during the first week of administration, but certain specific yet unidentified niche factors suppressed early B cell IgE switching in SLOs.

**Figure 2.**
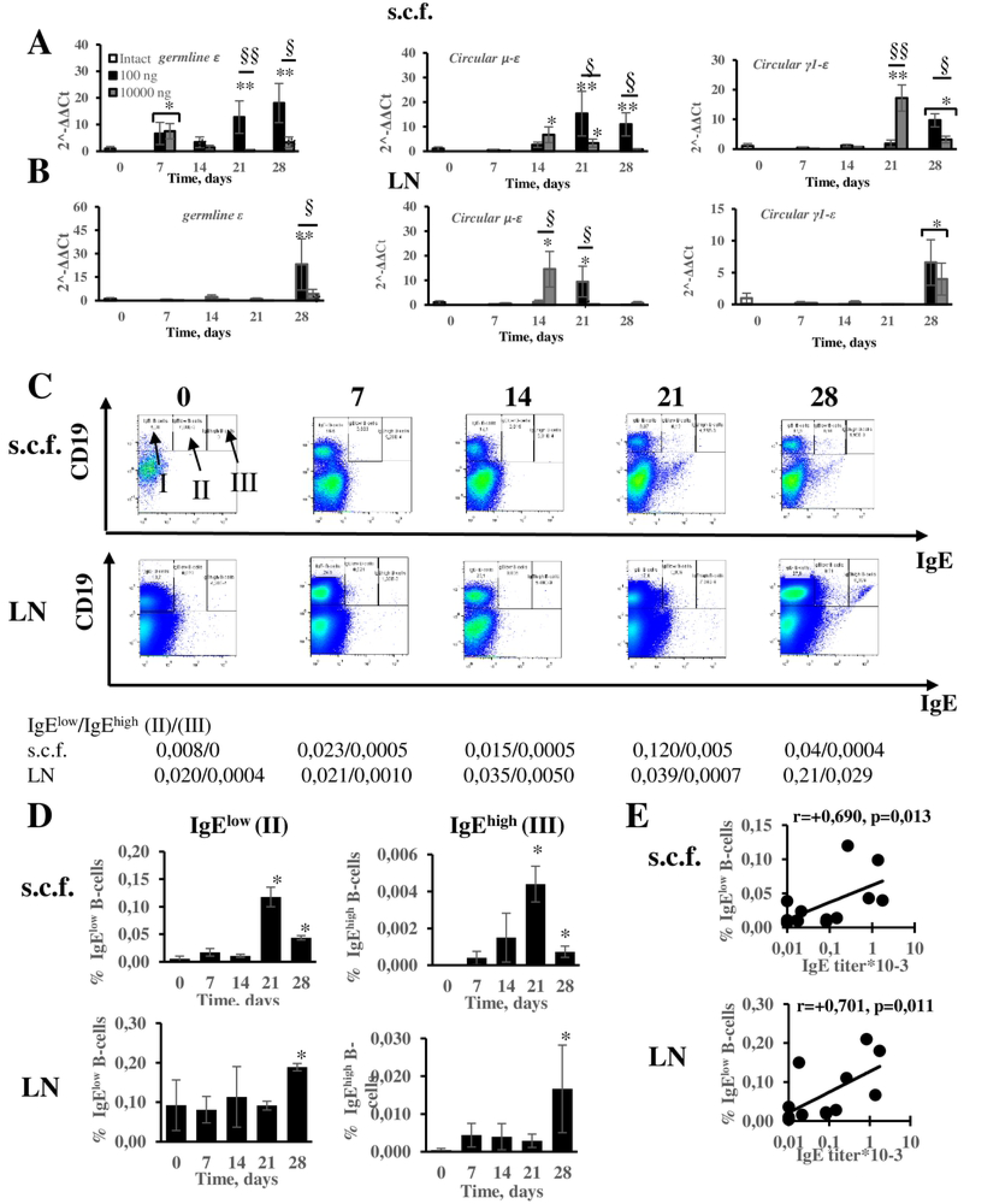
B-cell IgE class switching occurs in withers adipose tissue prior regional lymph nodes and leads to the formation of IgE^low^ and IgE^high^ B-cells. Both adipose tissue and lymph node IgE^low^ B-cells are associated with specific IgE production. Expression of germline transcripts and presence of DNA excision circles linked with IgE class switching in subcutaneous withers fat tissue (s.c.f.) (A) and regional lymph nodes (LN) (B). Representative flow cytometry pseudocolor plots (C). Roman numbers corresponds to following subpopulations: I – IgE- B-cells; II – IgE^low^ B-cells; III – IgE^high^ B-cells. Presence of IgE^low^ and IgE^high^ B-cells in withers tissue and lymph nodes (D). Correlations between IgE^+^ cells content in withers tissue or lymph node lymphocytes and IgE production (n=12) (E). */** - p <0.05/0.01 between indicated group and intact mice. §/§§ - p<0.05/0.01 between crossbars marked groups.

To investigate the relative contribution of IgE^low^ and IgE^high^ B-cells from tissue and lymph nodes in IgE production, we verified the presence of correlations between IgE production and percentage of these cells in tissue or lymph node respectively. Surprisingly, we have found that the quantity of IgE^low^, but not IgE^high^, B-cells is linked with specific IgE production. Despite that initial IgE class switching occurred in the site of antigen administration in tissue, the lymph node B-cells also participated in IgE production (Fig. 2E). There were no significant correlations between IgE production and quantity of IgE^high^ B-cells either in tissue or lymph nodes (Fig. S2). One could suppose that there are at least two pathways of IgE^+^ cells generation in our model. The first one leads to generation of B-cells which express high IgE^+^ BCR levels but do not differentiate effectively in IgE^+^ plasma cells. The second pathway leads to generation of B-cell express low levels of IgE^+^ BCR but rapidly differentiate into IgE^+^ plasma cells.

Two types of DNA excision circles corresponding to direct (µ-ε) or sequential (γ1-ε) IgE isotype switch appeared in our model. The excision circles corresponding to direct switch appeared, in general, earlier in both withers tissue and lymph nodes (Fig. 2A-B). We also have shown that quantity of IgE^low^ B-cells (but not IgE^high^) correlated with germline ε transcript expression and appearance of µ-ε DNA excision circle (Fig. S3). So, it is more likely that early IgE production is dependent on direct IgE switch from IgM to IgE which is consistent with our previous data obtained from young allergic patients [20].

It is well known that visceral adipose tissue contains a large number of FALCs [12]. FALCs resembling structures were found in human subcutaneous adipose tissue [13]. In present study to visualize these structures, we performed H&E histological staining of subcutaneous fat from withers region. Fig. S4 clearly shows that some fat-associated B-cells formed FALCs- like structures of different sizes, the others tended to form small dense or large diffuse infiltrates between adipocytes. Apparently these infiltrates could later develop to FALCs. This fact may indicate that FALCs in subcutaneous fat are very dynamic structures. This hypothesis is indirectly proven by different FALCs sizes as well as by communication of larger FALCs with lymphatic vessels (Fig. S5).

### IgG1 class switching occurs simultaneously both in subcutaneous fat-associated B-cells and lymph node B-cells

In contrast to IgE, IgG1 class switching was initiated simultaneously both in subcutaneous adipose tissue and regional lymph nodes (Fig. 3A) as characterized by germline γ1 expression. It is interesting that its expression was upregulated only at 21^th^ day at low dose group and at 28^th^ day at high dose group while IgG1 production appeared after 2 weeks of antigen adminstration. Apparently, low levels of B cells’ co-stimulation by T cells and germline transcript induction, which was undetectable in bulk tissue sample, were sufficient for IgG1 class switch, in contrast to IgE [43]. Indeed IgG1^+^ B-cells accumulation could be detected in both adipose tissue lymphocyte pool and in regional lymph nodes (Fig. 3B-C). However, only lymph node IgG1^+^ B-cell quantity correlated with specific IgG1 production (Fig. 3D). So, IgG1^+^ production induced by low dose antigen administration occurred mainly in regional lymph nodes.

**Figure 3.**
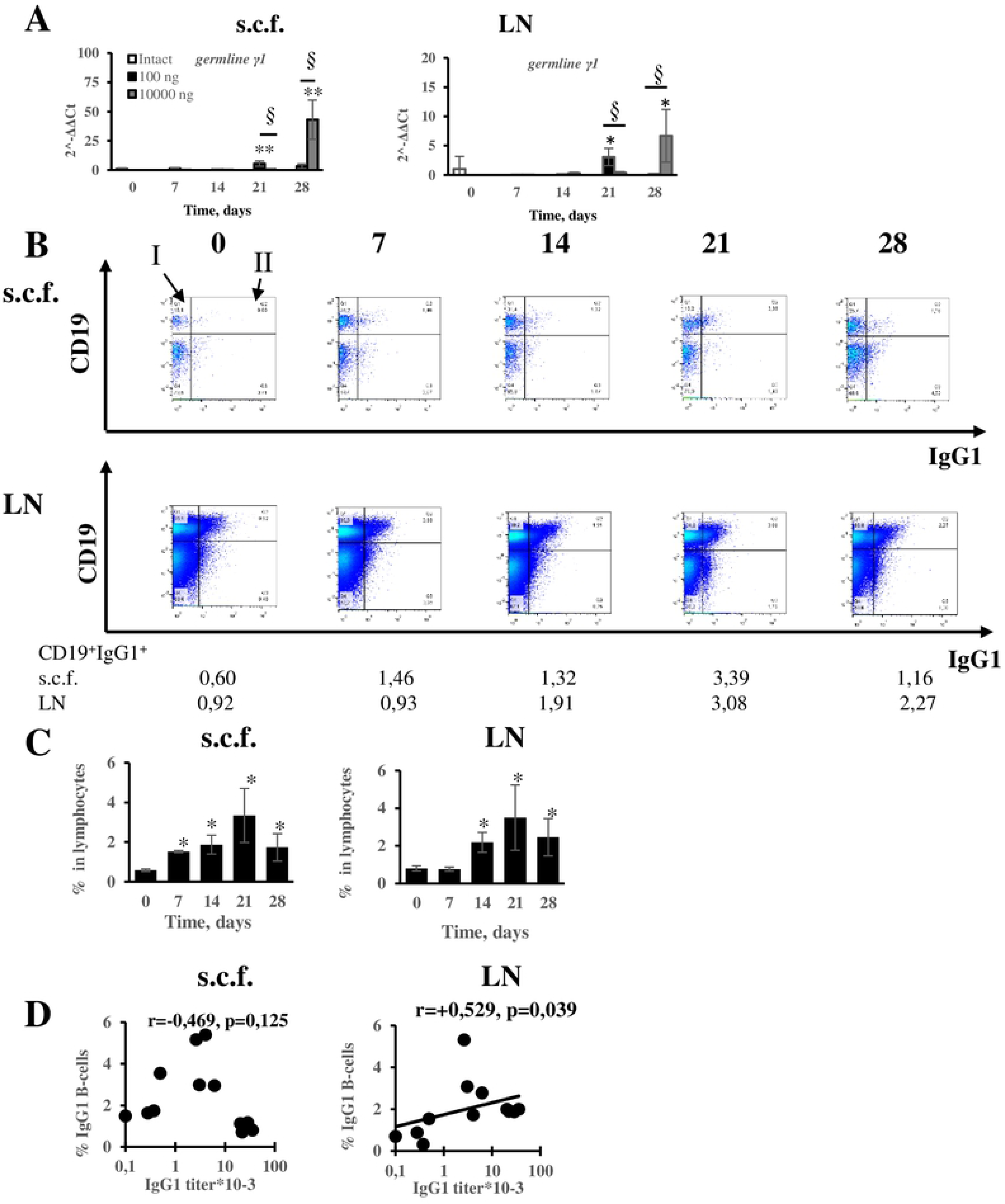
IgG1 class switching occurs at the same time points after the start of antigen administration in tissue and regional lymph nodes, but bulk levels of IgG1 antibodies are produced exclusively by IgG1^+^ B cells in regional lymph nodes. Expression of germline γ1 transcripts (A). Representative flow cytometry pseudocolor plots (B). Roman numbers corresponds to following suppopulations: I – IgG1- B-cells; II – IgE^low^ B-cells; III – IgE^high^ B-cells. Quantification of IgG1^+^ cells in subcutaneous fat withers tissue (s.c.f.) and regional lymph nodes (LN) (C). Correlation of IgG1^+^ cells’ relative numbers with IgG1 titers (D). */** p <0.05/0.01 between indicated group and intact mice. §/§§ - p<0.05/0.01 between crossbars marked groups.

### IgE^low^ B-cells acquire extrafollicular phenotype compared to IgE^high^ and IgG1^+^ B-cells

The tendency of IgElow B cells to develop into plasma cells more rapidly may be explained by the localization attachment of IgE low and IgE high populations within to different B cell compartments and undergoing separate differentiation pathways.. To investigate this possibility, we performed phenotype analysis of these cell subpopulations using B-cell markers which are differently expressed on naïve B-cells, GCs and plasmablasts – B220, CD38 and CD95. IgG1^+^ cells and total B-cell fraction were analyzed for comparison. Fig. 4 shows that at the beginning of IgE production (14^th^ day) all three B cell subpolulations with switched Ig isotype expressed significant levels of B220, though its expression on IgE^low^ cells was weaker compared to IgG1^+^ cells. Both IgE^low^ and IgE^high^ B-cells, in contrast to IgG1^+^ populations, were CD95 negative. Since B220 is a major B-2 cells marker except plasmablasts [34, 36], high levels of CD38 and CD95 indicates naïve and GC forming activated B-cells, respectively [35, 44] our data suggest that IgG1^+^ B-cells isotype switching, compared to IgE switching, occurs in cells that are more predisposed to GC formation.

**Figure 4.**
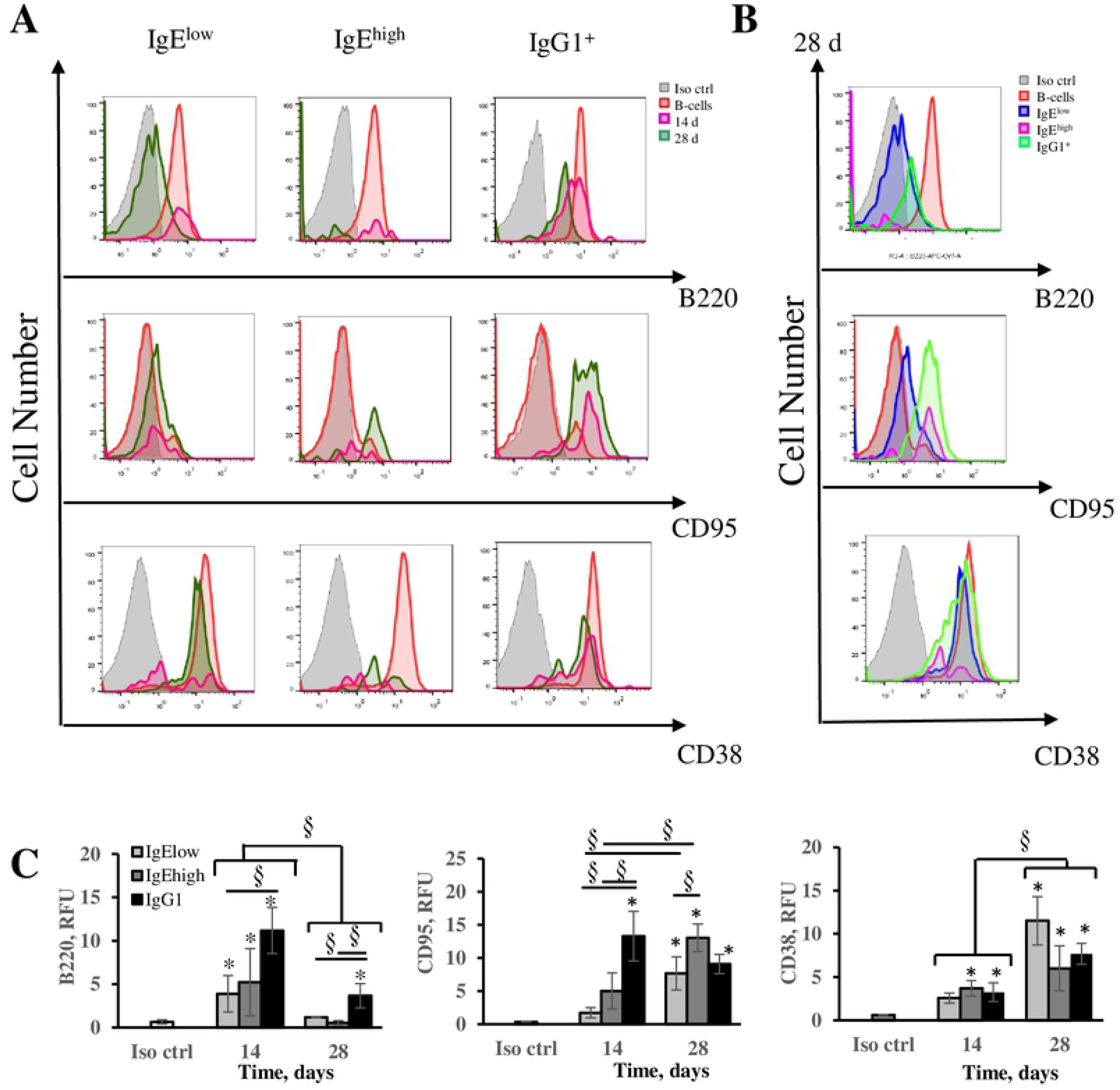
Acquisition of plamablastic phenotype by IgE^low^ rather than IgE^high^ and IgG1^+^ B-cells. Expression of B220, CD95 and CD38 on IgE^low^, IgE^high^ and IgG1^+^ B-cells. Representative flow cytometry histograms comparing isotype control (thin line, grey filled), total B-cells fraction (medium line, red filled), and respective subcutaneous fat tissue B-cell subpopulation on 14^th^ day of (thick line, crimson) or on 28^th^ day of OVA administration (thick line, dark green) (A). Representative histograms comparing to isotype control (thin line,grey filled), withers’ B-cells fraction (medium line, red filled), IgE^low^ B-cells (thick line, dark blue), IgE^high^ B-cells (thick line, pink), IgG1^+^ B-cells (thick line, light green) on 28^th^ day (B), Histograms showing relative B220, CD95 and CD38 expression (C). */** - p <0.05/0.01 between indicated group and RFU of isotype ctrl labeled sample. §/§§ - p<0.05/0.01 between crossbars marked groups.

However, at final time point (4 weeks, 28^th^ day) the expression of B220 was significantly decreased in all switched B-cell subpopulations IgE^low^ and IgE^high^ B-cells were completely negative, while IgG1^+^ cells still remained B220 positive in relevance to appropriate isotype control staining. Both IgE^low^ and IgE^high^ B-cells acquired substantial CD95 expression but IgE^high^ B-cells became almost all CD95 positive, whereas significant fraction of IgE^low^ cells remained CD95 negative. Mean expression of CD38 tended to increase in all subpopulations. But in the case of IgE^high^ and, to lesser extent, IgG1^+^ populations, there was significant fraction of cells with low to negative expression of CD38 (Fig. 4). These facts indicate that IgE^low^ B-cells, in contrast to IgE^high^ and IgG1^+^, tended more to develop into extrafollicular B-cell plasmablasts. As far as IgE^+^ cells are committed to programmed cell death due to high pro-apoptotic potential in GCs [45], the pathway which led to plasmablasts formation could support IgE response more actively than conventional activation which led to GC formation. We presume that separate B cell compartments were differentiated by similar though slightly different pathways and these discrepancies could be linked to different capacity to develop into CD19^+^B220^-/low^ plasmablast compartment Different rate of plasmablasts accumulation could results in different rate and intensity of IgE class switching in subcutaneous adipose tissue compared to regional lymph nodes.

### B-2 cell derived plasmablasts but not GCs are responsible for IgE production after long-term antigen administration

To verify this hypothesis, we performed flow cytometry analysis of B-cell subpopulations isolated from withers and regional lymph nodes at different time points during long-term antigen administration protocol in subcutaneous adipose tissue.

B-cell gating strategy is shown on Fig. S5. Fig. 5 clearly shows that there was no GCs (CD19^+^B220^+^CD38^-^CD95^+^) induction either in subcutaneous withers adipose tissue or in regional lymph nodes. Although high doses of antigen induced GC B-cell accumulation, in subcutaneous fat this induction was transient and was detected only at 21^th^ day, which indicate that conditions for GC persistence in subcutaneous fat were unfavorable, and in regional lymph nodes this induction was detected only at 28^th^ day. Instead, significant accumulation of CD19^+^B220^-^ plasmablasts was observed. Most of these plasmablasts were CD19^+^B220^-^CD38^-^CD95^+^ (Fig. 5C-D) and the amount of these cells in subcutaneous fat or lymph nodes directly correlated with specific IgE production (Fig. 5C, E). The absence of CD38 and presence of CD95 may indicate that these cells are closely relative to classical GCs differing from that only by the absence of B220 expression. The other possibility is that this phenotype could simply reflect full activated B-cell state.

**Figure 5.**
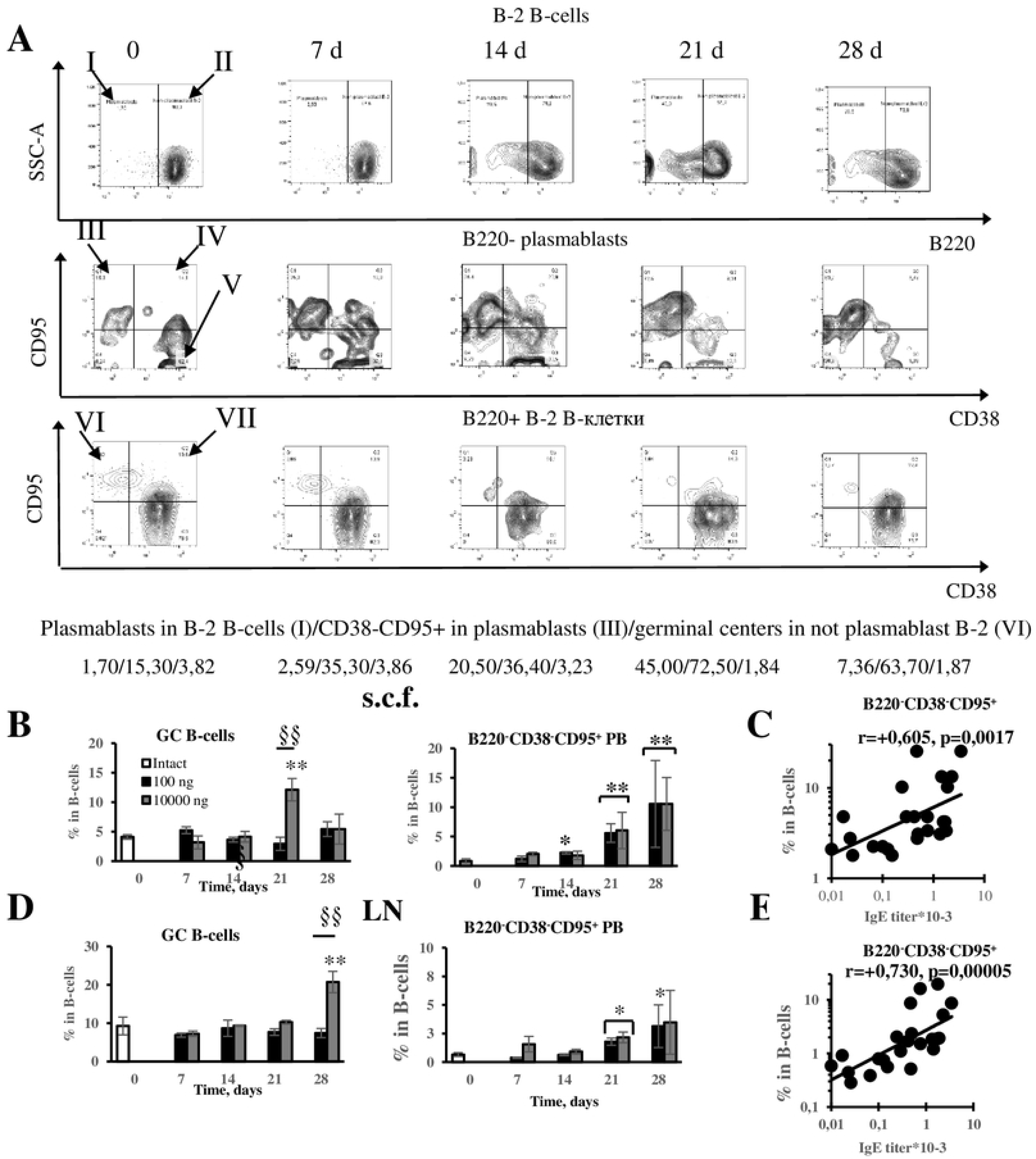
B220- plasmablasts derived from B-2 B-cells, but not GCs, are induced in withers and regional lymph nodes by low antigen dose and responsible for specific IgE production. Representative flow cytometry contour plots of indicated withers B-cells subpopulations at different time points (A). Roman numbers corresponds to the following subpopulations: I – plasmablasts, II – Not plasmablast follicular B-2 B-cells; III – CD38^-^CD95^+^ plasmablasts; IV – CD38^+^CD95^+^ plasmablasts; V – CD38^+^CD95^-^ plasmablasts; VI – germinal centers; VII – CD38^+^CD95^+^ not plasmablast B-2 B-cells. Relative amount of GC B-cells and B220^-^CD38^-^CD95^+^ plasmablasts in all CD19^+^ B-cells from withers tissue (B) and regional lymph nodes (D). Correlations of relative B220^-^CD38^-^CD95^+^ plasmablasts amount in subcutaneous withers fat tissue (s.c.f.) (C) and regional lymph nodes (LN) (E) with specific IgE titers (n=24). */** - p <0.05/0.01 between indicated group and intact mice. §/§§ - p<0.05/0.01 between crossbars marked groups.

The other plasmablasts subpopulations also accumulated in adipose tissue and regional lymph nodes and the earlier subpopulation in adipose tissue apparently represents B220^-^ CD38^+^CD95^-^ resembles naïve B-cells (Fig. S6A-B). It is likely that these cells later differentiate into other plasmablast subpopulations. The percentage of all of these subpopulations in CD19^+^ B-cells, however, did not significantly correlate with IgE production with exception of CD19^+^B220^-^ CD38^+^CD95^+^ and СD19^+^B220^-^CD38^-^CD95^-^cells in regional lymph nodes (Fig. S6C-D). As shown above, IgE switched B-cells acquired CD95 and CD38 expression over time and lost B220 expression. So, it is likely that after IgE class switching CD19^+^B220^-^CD38^+^CD95^+^ cells could differentiate from CD19^+^B220^-^CD38^-^CD95^+^ cells and CD19^+^B220^-^CD38^-^CD95^-^ from CD19^+^B220^+^CD38^-^CD95^-^, respectively. It is also interesting, that at early time points we could not detect significant differences in accumulation among various plasmablasts subpopulations, except CD19^+^B220^-^CD38^+^CD95^-^ in lymph nodes, in low and high dose immunized mice.

The percentage of GC B-cells in withers adipose tissue inversely correlated with specific IgE production. There was no functional association between the number GC B-cells in local lymph nodes and IgE levels (Fig. S7). As seen in gating strategy plots (Fig. S5), these plasmablasts subpopulations were not derived from B-1a or MZ-B B-cells. In comparison to B-2 derived plasmablasts no increase in B-1a or MZ-B B-cells was detected before day 21 upon immunization. Only at later time points there was transient increase in amount of B-1a B-cells (Fig. S8). However, this increase did not correlate with IgE production (data not shown).

In addition, we observed that the percentage of different plasmablasts subpopulations in regional lymph nodes and withers tissue started to increase at the same time points. It means that antigen even at low doses is rapidly delivered in regional lymph nodes. So, one can suppose that extrafollicular plasmablasts accumulation *per se* is necessary but not sufficient for IgE production, and the impact of different types of T-helpers on these cells in withers adipose tissue and regional lymph nodes could result in delayed IgE switching in regional lymph nodes.

### Extrafollicular T-helper cells accumulation results in high IgE production which is accompanied by minimal IgG1 production in response to low antigen doses

Indeed, different effector cytokine-producing cells and T-helper subpopulations could be responsible for specific humoral response pattern in response to low antigen dose. Different activity of these cells in withers tissue and regional lymph nodes could account for delayed IgE isotype switching in regional lymph nodes vs. subcutaneous adipose tissue B-cells.

Gating strategy for T-helper cell subsets, NK-cells and ILC2 cells is shown of Fig. S9. Presuming that extrafolliculary proliferating plasmablasts account for specific IgE production, it is logical that this production is also directly linked with extrafollicular T- helpers which are specific T-helper cell type supporting extrafollicular plasmablasts [46]. The IgE production is reciprocately linked with T-follicular helpers which support GC function and suppress the exit of B-cells from GCs by stimulating Bcl-6 expression [47]. Indeed, we observed remarkable decrease of CXCR4^+^CXCR5^+^ GC T-follicular helpers and CXCR4^-^CXCR5^+^ СD4^+^ T-helpers in subcutaneous fat tissue on 21^th^ day when IgE response had reached plateau. We suggest that CXCR4^-^CXCR5^+^CD4^+^ cells could be T-helpers that resided in B-cell follicles outside of GCs and these cells could also support B-follicle structure [38]. Decreasing of these T-helper cells subpopulations could account for GCs and B-cell follicles destabilization and competitive enhanced development of extrafollicular plasmablasts (Fig. 6). In contrast, CXCR4+CXCR5- T- extrafollicular helpers accumulated in subcutaneous fat tissue though only in low dose immunized mice and transiently on 21^th^ day. There was no accumulation of these cells in regional lymph nodes, whereas CXCR4-CXCR5+ T-helpers that support B-cell follicles accumulated significantly, particularly, in high dose group. Accumulation of GC supporting CXCR4^+^CXCR5^+^ T-follicular helpers was also seen in regional lymph nodes at 21th day in high dose group (Fig. 6). So, decreasing of CXCR4^+^CXCR5^+^ and CXCR4^-^CXCR5^+^ T-helpers activity percentage in CD4^+^ T-cells, which stabilize GCs, and B-cell follicles, respectively, results in accelerated specific IgE production in subcutaneous fat. CXCR4^+^CXCR5^-^ extrafollicular T- helpers accumulation could support specific IgE but not specific IgG1 production after long-term antigen administration at the levels compared to high dose immunized mice.

**Figure 6.**
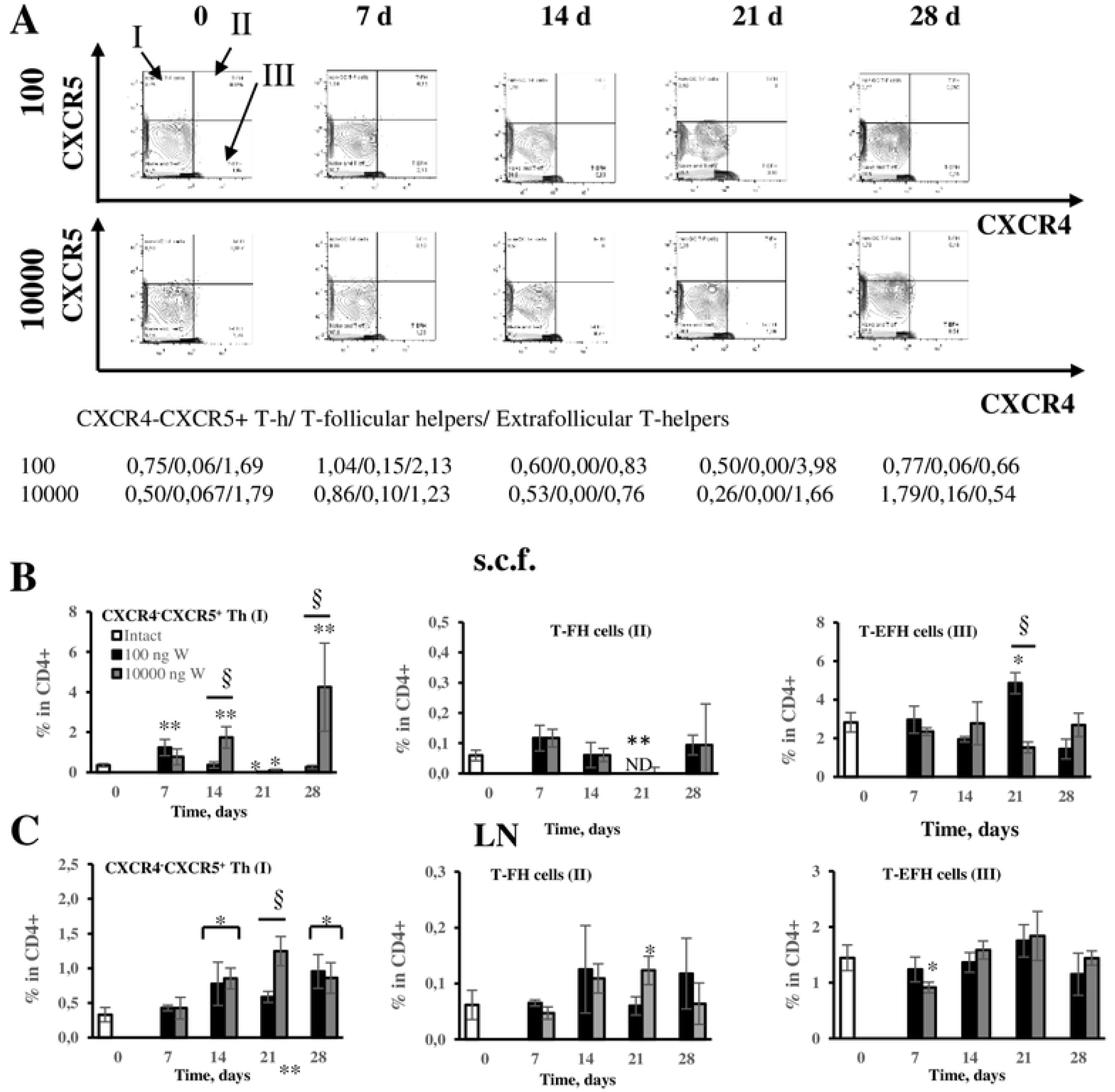
Extrafollicular T-helper cells, rather than follicular T-helpers or CXCR4^-^CXCR5^+^ T- helpers, are responsible for the formation of significant IgE production with minimal IgG1 response in withers adipose tissue after low dose antigen administration. Representative flow cytometry contour plots of T-helpers subpopulations in subcutaneous withers fat tissue of low and high dose immunized mice at different time points (A). Roman numbers corresponds to the following subpopulations: I – CXCR4^-^CXCR5^+^ T-helpers; II – CXCR4^+^CXCR5^+^ T-follicular helpers; III – CXCR4^+^CXCR5^-^ extrafollicular T-helpers. Quantification of T-helpers subpopulations in subcutaneous withers fat tissue (s.c.f.) (B) and regional lymph nodes (LN) (C). */** - p <0.05/0.01 between indicated group and intact mice. §/§§ - p<0.05/0.01 between crossbars marked groups.

It is also interesting that we did not observe significant accumulation of classical T-helper 2 cells (CXCR4^-^CXCR5^-^ST2^+^) which resided mainly in T-cell zone [39] after low dose antigen administration. Even high antigen doses induce their accumulation only at 28^th^ day and mainly in subcutaneous adipose tissue but not in regional lymph nodes (Fig. S10). These cells could support later stages of IgE production.

ILC2 cells are usually mentioned in context of immune reaction to tissue damage and sterile inflammation [48]. Although ILC2 showed some tendency to accumulate in subcutaneous fat, they were too rare and this tendency was insignificant (Fig. S11). Still we observed accumulation of CD4^-^ and CD4^+^ NK^-^cells (CD49b^+^) at the end of immunization in subcutaneous fat of high dose immunized mice and in regional lymph nodes in both groups (slightly earlier at low dose) (Fig. S11). Due to their late accumulation in the tissue, these subpopulations could not be associated with early IgE production.

### Delayed and hampered IgE and IgG1 B-cell formation in abdominal fat tissue after upon long-term antigen administration is due to unstable induction of extrafollicular B-cell plasmablast accumulation and absence of extrafollicular T-helpers accumulation

It is well known that abdominal (visceral) fat tissue contains large numbers of FALCs where local immune response can be initiated [11, 12]. Despite his fact long-term intraperitoneal antigen administration into the region enriched in visceral fat and associated FALCS established significant lower and delayed IgE and IgG1 production (Fig. 7 A-E). So, one can conclude that subcutaneous fat (in our case, in withers region) connected more closely to skin epidermis has markedly different properties in comparison to visceral abdominal fat which is not connected directly to skin barrier. The diversity in immune response in both analyzed fat tissues in low dose model may be functionally associated with differences in subcutaneous and visceral fat-associated immune cells.

**Figure 7.**
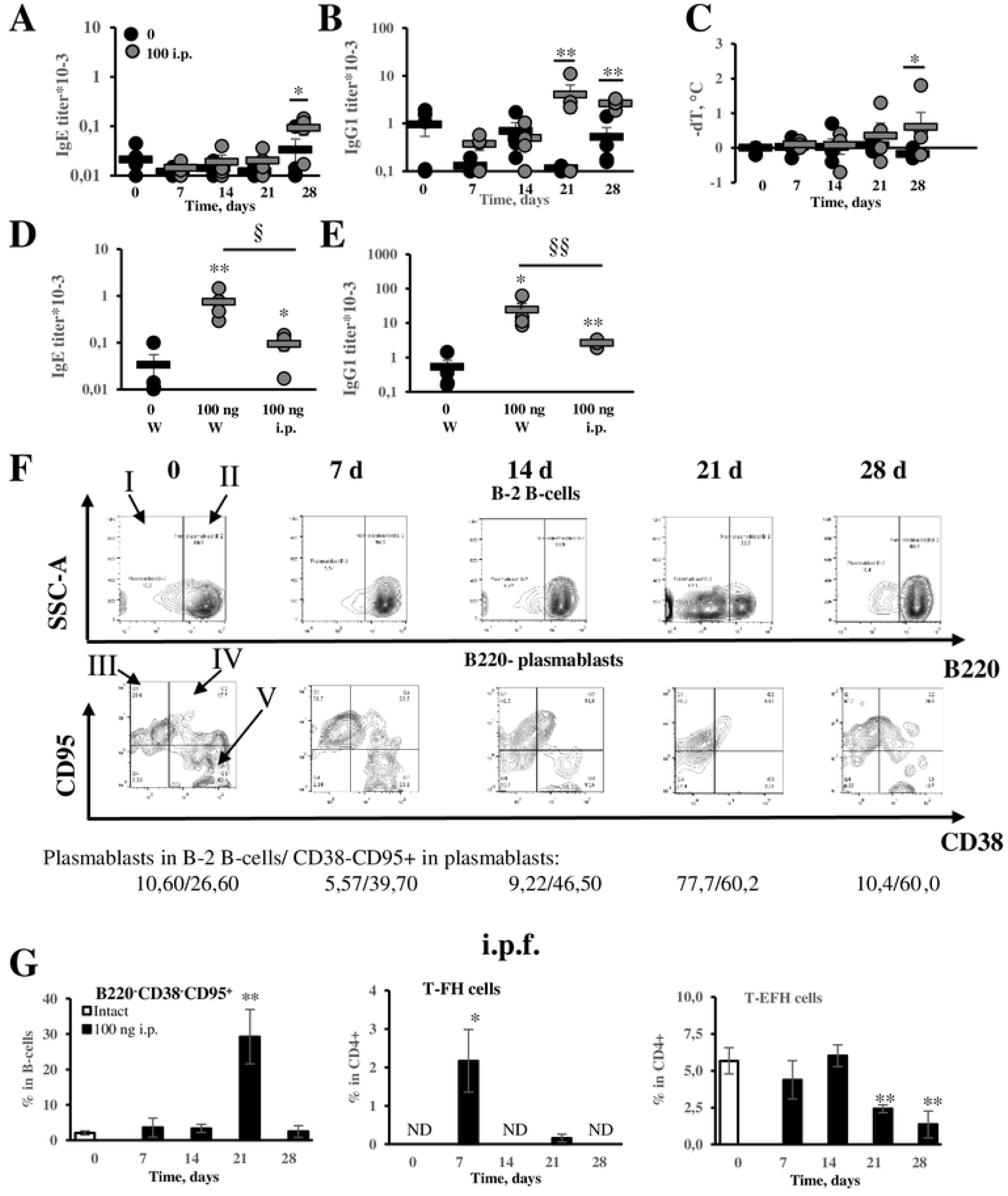
Transient and unstable induction of extrafollicular plasmablasts, and lack of extrafollicular Th accumulation are responsible for delayed and hampered antibodies production in abdominal fat. BALB/c mice were immunized by low OVA dose (100 ng) by i.p. route. Specific IgE (A), IgG1 (B) titers and anaphylactic severity (C) were measured and compared with respective parameters of saline immunized animals. Comparison of specific IgE (D) and IgG1 production (E) between mice immunized by low antigen dose via subcutaneous route in withers tissue and by i.p. low dose immunized mice. Representative flow cytometry plots of indicated cell populations in abdominal fat tissue (i.p.f.) at different time points (F). Roman numbers corresponds to the following subpopulations: I – plasmablasts; II – not plasmablast B-2 B-cells; III – CD38^-^CD95^+^ plasmablasts; IV - CD38^+^CD95^+^ plasmablasts; V – CD38^+^CD95^-^ plasmablasts. Percentage of indicated cell subpopulations (G). */** - p <0.05/0.01 between indicated group and intact mice. §/§§ - p<0.05/0.01 between crossbars marked groups.

To understand the mechanism of this phenomenon, we compared B- and T-cell activation in subcutaneous and abdominal fat tissues upon antigen administration in low doses model. The low concentrations of antigen were administered by either subcutaneous injection in withers or intraperitoneally according to 28 days immunization protocol. First, we wondered if delayed kinetics of IgE and IgG1 formation in abdominal FALCs could be due to delayed appearance of IgE^+^ and IgG1^+^ B-cells Indeed, we observed that upregulation of germline ε transcripts in abdominal fat occurred later than in subcutaneous fat tissue (Fig. S12 A-B). After day 21 of antigen administration IgE^+^ B cells started to accumulate in subcutaneous fat and reach plateau, presumably, due to rapid differentiation of CD19^+^ IgE-producing plasma cells by day 28. The percentage of IgE+ cells in abdominal fat was lower, which indicated more delayed kinetics of B cells accumulation in visceral adipose tissue. (Fig. S12 C-D). The germline γ1 transcripts were upregulated in both adipose tissues at the same time point. In abdominal fat, their induction was transient and diminished to initial level by 28^th^ day (Fig. S12 A-B). IgG1^+^ cells accumulated more rapidly in subcutaneous fat than in abdominal (Fig. S12 D).

Delayed IgE and IgG1 production in abdominal fat tissue could be caused by reduced of extrafollicular B-cell activation in this site compared to subcutaneous fat tissue. Indeed, in abdominal fat tissue, the accumulation of CD19^+^B220^-^CD38^-^CD95^+^ plasmablasts which were responsible for IgE production in subcutaneous fat, was transient and unstable (Fig. 7 F-G). Such a weak plasmablasts activation can be explained, firstly, by early burst of Tfh activity in abdominal fat tissue on day 7 upon of long-term antigen administration, and, secondly by decrease in extrafollicular Th amount at later time points (3-4 weeks of the immunization) (Fig. 7 G and Fig. S13). We did not observe significant GC B-cells accumulation in abdominal fat which could be due to absence of CXCR4^-^CXCR5^+^ T-helpers which function was to stabilize B-cell follicles (Fig. S14). So, early Tfh burst only could hamper extrafollicular B-cell differentiation instead of supporting normal GC B-cell development The activity of extrafollicular T-helpers is essential for stable Ig production by extrafollicular B cell plasmablasts during low dose immunization regime when low antigen concentrations cannot initiate strong BCR-mediated signaling. We presume that in the absence of extrafollicular T helpers activity, the accumulation of plasmablasts could be transient and did not cause strong antibody production.

After second week of immunization, CD4^+^ NK cells started to accumulate in abdominal fat. Minor increase in CD4^-^ NK-cells was detectable after 7^th^ day of immunization (Fig. S15). CD4^+^ NK-cells and its CD1d-restricted variant CD4^+^ iNKT cells could activate [49] or inhibit B-cell Ig production [50] depending on special conditions. The potential inhibitory role of NK cells is an aim of further studies.

## Discussion

In this study, we found the site of antigen-induced B cell IgE isotype switching in low doses allergen model (1) and investigated the main patterns of B- and T-cell activation. In our allergy model, the route of immunization and allergen dose plays essential role in humoral immune response. Although long term immunization with both low and high antigen doses induced comparable levels of IgE production, the kinetics of IgE production upon high dose immunization was slightly delayed and reached plateau only after day 28 of immunization compared to day 21 of immunization with low dose. Moreover, only in low doses administration protocol, anaphylaxis severity correlated with specific IgE production which resembles clinical manifestations in humans [20; 51; 52]. Our data are in agreement with previous research reporting that long-term low dose allergy models better reflect natural sensitization process [24–27]. Both subcutaneous fat-associated and local lymph node B cells were activated upon long term antigen administration and produced specific IgE antibodies. Subcutaneous fat was the primary site of antigen-induced IgE class switching This finding is important for further development of novel approaches of allergen-specific immunotherapy aiming to prevent IgE class switching and eliminate both local and systemic IgE-producing B cells.

After long-term antigen challenge two IgE^+^ B-cells subpopulations were clearly observed in contrast to IgG1^+^ cells. These cells arose from pools of B cells which underwent distinct diffrentiation pathways following isotype switching. The first subpopulation, IgE^low^ cells, seemed to be closely linked with IgE production in contrast to other pool, IgE^high^ B-cells. In general, IgE^+^ cells more efficiently differentiated into plasmablasts compared to IgG ^+^ cells (as judged by loss of B220 expression). In contrast to IgE^high^, IgE^low^ B cells acquire phenotype more distinct from the phenotype of activated B cells resembling GCs which is characterized by low CD95 and higher CD38 expression. Although we could not clearly resolve this question we suppose that IgE^low^ cells are more predisposed to develop into plasmoblasts meanwhile IgE^+^ cells with GC phenotype are more prone to apoptosis [45] unless they rapidly exit GC and differentiate into antibody producing cells which do not give rise to stable IgE memory formation [53]. The observed differences in CD95 expression levels on IgE^low^ and IgE^high^ cells are of particular interest because of essential role of CD95 molecule in eliminating self-reactive and IgE^+^ cells [54]. Although conventional plasmablasts are not believed to form GC-like structures, but in present study, we have shown that some B220^-^ plasmablasts acquired GC-like phenotype (CD38^-^CD95^+^) and precisely these cells are engaged in specific IgE production. Recent research works showed that plasmablasts could also be regulated by extrafollicular T- helpers, and this regulation is crucial for long-term Ig production [38, 46]. These Th cells closely interact with extrafollicular B-cells. While CD95 expression on murine B-cells is induced by CD40 ligation [44], and CD38 expression is inhibited upon T-dependent GC formation, one can conclude that at certain circumstances extrafollicular B-cells also acquire GC-like phenotype. Later, immediately before differentiation into Ig producing cells, these cells, especially IgE^low^, change their phenotype and lose GC-like properties. This can be due to weaker B-T cell contacts on IgE^low^ cells which express low BCR levels and, therefore, cannot present antigen as efficiently as cells expressing high BCR levels. [55]. Despite that IgE^+^ cells mostly acquired CD38 expression, the closer correlation between IgE production and B220^-^CD38^-^ CD95^+^cells rather than B220 plasmablasts was observed. One possible explanation is low probability of cognate B-T cell contacts when low doses of antigens are administrated. The stage when extrafollicular B cells form these contacts and acquire GC-like phenotype is pivotal for the whole process of IgE-producing cells formation. The relations of IgE production and B220^-^ CD38^+^CD95^+^ cells accumulation in lymph nodes were significant and more prominent, in comparison to that seen in adipose tissue. The lymph nodes possess special stromal cell architecture which provides the more optimal niche for B cell proliferation and survival compared to non-lymphoid tissues. It is tempting to assume that B220^-^CD38^+^CD95^+^ cells represent the final stage of IgE^+^ cells differentiation into plasma cells before they lose CD19 expression. Further investigations are needed to clarify this hypothesis.

If B220^-^CD38^-^CD95^+^ plasmablasts are cells that contact with T cells during response to low antigen doses, it can be assumed that the presence of such Th subpopulation is crucial for development of IgE response. Indeed, we show here that accumulation of extrafollicular T- helper cells occurs, first, only in subcutaneous fat tissue where the early, initial IgE class switching is induced rather than in regional lymph nodes, and, second, only in low dose immunized mice. Despite being transient and visible only 3 weeks after the start of immunization, this accumulation is appeared to be critical for maintaining high levels of IgE production. On the contrary, despite early B cell differentiation into plasmablasts in regional lymph nodes as a result of quick antigen delivery, IgE class switch was not detected. Moreover, after 3-4 weeks upon antigen administration into abdominal fat tissue, we observed a decrease in percentage of extrafollicular Th cells. This seems to be a plausible reason for transitory but not stable B220^-^CD38^-^CD95^+^ plasmablasts accumulation in these sites and subsequent delayed IgE and IgG1 production. Besides, in subcutaneous fat, we observe marked decrease in percentage of Tfh and CXCR4^-^CXCR5^+^ Th in that occurred simultaneously when accumulation of extrafollicular T- helper and IgE response reached plateau. In the absence of these subpopulations that support B-cells in GCs [47] and mantle zone [38], B-cell follicular response can be dropped down and extrafollicular B-cell activation competitively increases. High antigen doses may cause significant antigen accumulation in either tissue or lymph nodes which results in elevated signal from BCR which alone could sustain B-cell proliferation and antibody production, as it was found by some research teams [8, 9]. So just response to low but not high antigen doses is linked with T-extrafollicular helpers accumulation.

The question why these T-cell subpopulations are differently regulated in subcutaneous fat tissue vs regional lymph nodes and abdominal fat tissue is not yet resolved. Apparently certain myeloid cells such as DC [56] and macrophages [57, 58] are able to act as regulators of T cells functions in lymphoid and non lymphoid tissues. However, these particular issues are beyond the scope of present study and will be addressed in future.

In our study, we did not use antibody-based or small molecular inhibitors of GCs. It is not evident that IgE isotype switching per se occurs in early GC B-cells which emigrate from these structures soon afterwards. First, in our work, the majority of IgE^+^ cells remained CD95^-^ CD38^-^

^/low^ after beginning of isotype switch which happened 2 weeks after the immunization start. This means that despite becoming activated these cells did not acquire full GC phenotype. Second, we did not observe any increase in GC B-cells after low dose antigen administration either in tissue or in regional lymph nodes and did not detect any positive correlations between these cells’ accumulation and IgE production. Despite that both class switch DNA recombination and somatic hypermutation occur within GCs [59], several studies suggest that extrafollicular B-cell class switch recombination is also possible at least at early stages of plasmablasts differentiation [60; 61]. Furthermore, recent work clearly shown that B-cell class switch recombination at least in some cases occurs mostly in early stages of B-cell activation before differentiation into GC centroblasts or extrafollicular B-cell blasts and is dependent mostly on T-B cell contacts per se but not on GC formation. Only somatic hypermutation is linked exclusively with GCs [62]. These findings support our data showing that 2 weeks after start of immunization the early IgE^+^ cells acquired non-fully activated phenotype characterized by B220^+^CD95^-/low^ expression. Later, after week 4, they differentiated into fully activated plasmablasts (B220^-^CD95^+^).

The low dose administration induced the extrafollicular but not GC-associated antibody switching, and, therefore, turned to be not favorable for somatic hypermutation, which is essential for high affinity antibody production. Somatic hypermutation always occurs within GCs [62]. These high affinity IgE are associated with severe anaphylactic reactions. However, we have clearly observed significant anaphylaxis response in mice (*Fig. 1*) This was in agreement with previously published data [22]. The possible explanation is that some occasional B-cells express surface BCR with relatively high affinity to novel antigen even prior to somatic hypermutation, and these particular cells differentiates into plasmablasts [63]. Our findings in are not consistent with recently published study which clearly demonstrated the essential role of IL-4^+^IL-13^+^ Tfh in production of high affinity IgE [64]. However, the authors used high dose immunization protocol in knockout mice with conditional deletion of DOCK8 in CD4^+^ T cells. In this model, DOCK8 deficiency reveals unique IL-4^+^IL-13^+^ Th set not only in T-follicular but also in T-extrafollicular cell compartment. Therefore, DOCK8 knockout could be responsible for increased IgE production not only due to appearance of IL-4-IL-13 double producers in T-follicular helpers but also in extrafollicular T-helpers [64]. Our results are in agreement with several clinical investigations which maintain that local B-cell IgE class switching mostly depend on extrafollicular B-cell activation [10, 65]. Earlier studies based on IgE reporter mice and adoptive transfer experiments of IgG1 switched B-cells suggest two waves of IgE^+^ cells generation. During the first wave extrafollicular response and direct µ-ε class switching were detected. The second wave is characterized by sequential γ1-ε class switching and GC response and gives rise to stable IgE production [66]. Our observation that IgE class switch occurs in extrafollicular B cells (Fig. 4, 5) partially confirms this theory.

We emphasize here that all plasmablasts in our model were derived from conventional B- 2 B-cells as verified by the absence of B-1a and MZ-B markers expression (Figure S5). We did not observe significant sustained accumulation of either B-1a or MZ-B cells. The participation of CD5-B220-CD19+ B-1b cells in extrafollicular IgE production is highly unlikely since these subsets rarely produce high or medium affinity antibodies and are predisposed to IgG and IgA class switching [68]. So, our extrafollicular IgE production must be T-cell dependent.

In our work, we have clearly shown that sustained plasmablasts activation is linked with extrafollicular T- helpers accumulation and rapid IgE class switching in subcutaneous adipose tissue in comparison to abdominal fat and regional lymph nodes. We could not completely identify cell populations which produced type 2 cytokines, such as IL-4, for the switching *per se* at the early stages. The current data did not provide evidence for the contribution of Th2 (Fig. S10). The tendency of ILC2 to accumulate in subcutaneous fat was insignificant due to high mouse-to-mouse variability. Meanwhile, the accumulation of NK cells as potential regulators of IgE response [50] in lymph nodes was detected only at later stages of IgE production, in relation to delayed IgE production in abdominal fat, which is in line with previous publication [50]. Further works will address these questions.

One of the most importance output from our work is that in allergically predisposed subjects, humoral immune response to low antigen doses entering due to defects in barrier tissues is markedly different in subcutaneous vs. abdominal fat. Numerous works are aimed to understand mechanisms of formation and function of fat-associated lymphoid clusters in visceral, usually abdominal, adipose tissue [69]. Despite that allergens penetrating skin barrier, in fact, enter the body through subcutaneous fat, a few studies [13, 14] yet provide information on fat-associated B-cell. Frasca et al. [14] observed that pro-inflammatory cytokines secreted by adipocytes promoted T-bet and CD11c expression on subcutaneous withers associated B-cells. The expression of these molecules on CD19^+^ B-cells identifies extrafollicular plasmablasts subpopulation with specific properties. These B-cells associated with subcutaneous adipose tissue are capable to form FALCs of different sizes as well as diffuse infiltrates [13, 70–72]. We can hypothesize that tissue variations in T-bet regulation may also contribute to remarkable differences in B and T cell response within subcutaneous vs. visceral fat. On the other hand, failures in extrafollicular response in visceral fat can be attributed to accumulation of different antigen presenting cells or regulatory CD4^+^NK cells within abdominal tissue upon antigen treatment (Fig. S15).

The transient nature of GCs formation in the regional lymph node requires particular explanations. We presume that certain adipocyte released mediators such as free fatty acids [73] may play a role by triggering unfolded protein response [74] in B cells resulting in enhanced plasma cells and plasmablasts development instead of stable GC persistence [75].

Overall, we show here that in subcutaneous adipose tissue, the response to low antigen doses is extrafollicular by its nature and based primarily on activation of CD19+B220-CD38-CD95+ plasmablasts. Accumulation of CXCR4^+^CXCR5^-^ extrafollicular T- help*e*rs is also critical for this response. This unique type of immune response creates specific pattern of humoral response which is characterized by high IgE production accompanied by minimal IgG1 production and leads to allergy development. Subcutaneous fat tissue but not abdominal fat tissue and SLOs provide the most favorable conditions for development of such response. Although some questions remains unresolved in our work, we suppose that these results are very important. It becomes clear that future development of new methods of allergy prevention in prone individuals should be aimed not only at stimulation of GC formation where protective IgG1 antibody-producing clones are formed [76] but also at extrafollicular B-cell response blockage. The latter in some cases may be even more important because in our model local environment in subcutaneous fat obviously supports extrafollicular but not GC response even at high dose regime of immunization. Novel potential therapeutics also must disrupt extrafollicular B-cell activation both in SLOs and in TLSs.

## Abbreviations

BCR: B-cell receptor
ELISA: Enzyme linked Immunosorbent Assay
FALCs: Fat associated lymphoid clusters
GC: germinal centers
iBALT: inducible bronchial associated lymphoid tissue
i.p.: intraperitoneal
LN: Lymph nodes
OVA: Ovalbumin
PCR: polymerase chain reaction
s.c.: subcutaneous
SLOs: secondary lymphoid organs
Tfh: T-follicular helpers
Th: T helpers
TLSs: tertiary lymphoid structures

## Acknowledgements

We thank Senior Scientist of Laboratory of Molecular Diagnostics IBCh RAS Ryasantsev D.Yu. for useful advices on primer and probe design and quantitative PCR performance. The reported study was founded by RFBR according to the research projects number 19-015- 00099 and 19-05-50064.

## Author contibution

Conceived and designed experiments: CDB, FGV., SEV. Performed experiments: CDB, FGV, KMV, TDS, SMA. Analyzed the data: CDB, KOD, SAA, SEV. Wrote the paper: CDB, FGV, SEV.

---

**Figure S1. Immunization protocol.** Black arrows indicate days of immunization. Red arrows indicate days of blood collection and mice sacrificing when samples of subcutaneous fat adipose tissue or abdominal fat tissue together with regional lymph nodes was taken.

**Figure S2. IgEhigh B-cells are not be responsible for specific IgE production**. Correlations between relative content of IgEhigh B-cells in subcutaneous withers fat tissue (s.c.f.) or lymph node (LN) lymphocytes and IgE production (n=12).

**Figure S3. IgE class switching in withers adipose tissue gives rise to the formation of IgE^low^, but not IgE^high^, B-cells, and occurs by direct mechanism.** Correlations between relative numbers of IgE^low^ (A-C) or IgE^high^ (D-F) B-cells in subcutaneous adipose tissue lymphocytes and expression of germline ε transcripts (A, D), relative quantity of circular µ-ε (B, E) or circular γ1-ε DNA excision fragments (n=12) (C, F).

**Figure S4. Subcutaneous fat adipose tissue in withers region contains tertiary lymphoid structures of different sizes as well as irregularly shaped lymphoid infiltrates.** Representative histological images of mouse subcutaneous withers adipose tissue. Tissue samples were taken from low dose immunized mice 4 weeks after the start of antigen administration. Samples were stained with hematoxilin – eosin and images were taken at 100X magnification. Adipose tissue is marked by white arrows, lymphoid clusters of different size are marked by black arrows. Dense lymphoid infiltrates characterized by irregular shape or presence of adipocytes are marked by grey arrows. Diffuse infiltrates are shown by brown arrows. Longitudinally cut lymphatic vessel associated with large lymphoid cluster is marked by blue arrows. The border of tertiary lymphoid structures and infiltrates is shown by red dashed lines.

**Figure S5. B-cells gating strategy.** Roman numbers corresponds to the following subpopulations: I - cells; II - Single cells; III – Live cells; IV – CD5^+^ B-cells; V – CD5^-^ B-cells; VI – B-1a cells; VII – MZ-B B-cells; VIII – Follicular B-2 B-cells; IX – Not plasmablast follicular B-2 B-cells; X – Plasmablasts; XI – CD38^-^CD95^+^ plasmablasts; XII – CD38^+^CD95^+^ plasmablasts; XIII – CD38^+^CD95^-^ plasmablasts; XV – germinal centers; XVI – CD38^+^CD95^+^ not plasmablast activated follicular B-2 B-cells.

**Figure S6. In both withers and lymph nodes, the antigen-dependent induction of other B220^-^ plasmoblasts populations is weaker than of B220^-^CD38^-^CD95^+^cells.** Presence of indicated plasmablasts subpopulations in withers adipose tissue (A) or regional lymph nodes (B) at different time points. Correlations of relative content of indicated plasmablasts subpopulations in subcutaneous fat tissue (s.c.f.) (C) or lymph nodes (LN) (D) with specific IgE titers. */** - p <0.05/0.01 between indicated group and intact mice. §/§§ - p<0.05/0.01 between crossbars marked groups.

**Figure S7. Percentage of GCs in withers adipose tissue and regional lymph nodes do not correlate with specific IgE production.** Correlations between relative B220^+^CD38^+^CD95^+^ GC B-cell numbers in withers fat tissue (s.c.f.) B-cells or lymph node (LN) B-cells (n=24).

**Figure S8. Long term antigen administration does not cause any stable increase in either B-1a or MZ- B cells in subcutaneous fat and regional lymph nodes.** Percentage of minor T-independent B-cell subpopulations in subcutaneous withers fat tissue (s.c.f.) (A) and regional lymph nodes (LN) (B) at different time points. */** - with p <0.05/0.01 between indicated group and intact mice.

**Figure S9. T-helper subsets, NK-cells and ILC2 cells gating strategy.** Roman numbers corresponds to the following subpopulations: I - cells; II - Single cells; III – Live cells; IV – Lin^-^CD45^+^ Cells; V – Lin^+^CD45^+^ cells; VI – ILC2; VII – T-helpers; VIII – CD4^+^ NK-cells; IX – CD4^-^ NK-cells; X – CXCR4^-^CXCR5^+^ cells; XI – T-follicular helpers; XII – Extrafollicular T-cells; XIII – Naïve T-cells and T-effectors; XIV – T-helper 2 cells.

**Figure S10. Low impact of Th2 cells on early stage of response to low antigen doses.** Representative flow cytometry contour plots (A) and percentage of ST2^+^CXCR4^-^CXCR5^-^ type 2 helper cells in total CD4^+^ Th fraction from subcutaneous withers fat tissue (s.c.f.) and regional lymph nodes (LN) of immunized mice in different time points (B). Roman numbers corresponds to the following subpopulations: I – T-helper 2 effector cells */** - p <0.05/0.01 between indicated group and intact mice. §/§§ with p<0.05/0.01 between crossbars marked groups.

**Figure S11. ILC2 and NK-cells do not participate in regulation of specific IgE production in withers adipose tissue but NK-cells potentially regulate in regional lymph nodes.** Representative flow cytometry images (A) and percents of indicated cell populations in subcutaneous withers fat tissue (s.c.f.) (B) or regional lymph node (LN) CD45^+^ cells (C). Roman numbers corresponds to the following subpopulations: I – CD4-CD49b^+^ NK-cells; II – CD4^+^CD49b^+^ NK-cells. */** - p <0.05/0.01 between indicated group and intact mice. §/§§ - p<0.05/0.01 between crossbars marked groups.

**Figure S12. Delayed kinetics of IgE class switching, IgE and IgG1 B-cell numbers increment in abdominal fat tissue in comparison to subcutaneous withers fat tissue after continuous low dose antigen administration.** Expression of indicated transcripts linked with Ig class switching in subcutaneous fat (s.c.f.) (A) and abdominal fat (i.p.f.) (B) in different time points during continuous antigen administration. Representative flow cytometry pseudocolour plots (C) and relative amounts of IgElow and IgG1^+^ cells in subcutaneous and abdominal fat tissue (D). Roman numbers corresponds to the following subpopulations: I – IgE- B-cells; II – IgE^low^ B-cells; III – IgE^high^ B-cells */** - with p <0.05/0.01 between indicated group and intact mice.

**Figure S13. Representative flow cytometry plots of T-helper cells subpopulations in abdominal fat tissue at different time points.** Roman numbers corresponds to the following subpopulations: I – CXCR4^-^CXCR5^+^ T-helpers; II – CXCR4^+^CXCR5^+^ T-follicular helpers; III – CXCR4^+^CXCR5^-^ extrafollicular T-helpers.

**Figure S14. No significant induction of GC B-cells and CXCR4^-^CXCR5^+^ T-helpers in abdominal fat tissue after low dose antigen administration.** Percentage of GC B-cells in B-cells and CXCR4^-^CXCR5^+^ T-cells in T-helpers of abdominal fat tissue at different time points. */** - with p <0.05/0.01 between indicated group and intact mice.

**Figure S15. CD4+, but not CD4-, NK-cells could probably regulate delayed humoral immune response in abdominal fat tissue.** Representative flow cytometry contour plots (A) and percent of CD4^-^ and CD4^+^ NK-cells (CD49b^+^) in CD45^+^ cells (B). Roman numbers corresponds to the following subpopulations: I – CD4^-^CD49b^+^ NK-cells; II – CD4^+^CD49b^+^ NK-cells. */** - p <0.05/0.01 between indicated group and intact mice. §/§§ - p<0.05/0.01 between crossbars marked groups.

